# H3K9 tri-methylation at *Nanog* times differentiation commitment and enables the acquisition of primitive endoderm fate

**DOI:** 10.1101/2021.06.22.449256

**Authors:** A. Dubois, L. Vincenti, A. Chervova, S. Vandormael-Pournin, M. Cohen-Tannoudji, P. Navarro

## Abstract

Mouse Embryonic Stem (ES) cells have an inherent propensity to explore distinct gene-regulatory states associated with either self-renewal or differentiation. This property is largely dependent on ERK activity, which promotes silencing of pluripotency genes, most notably of the transcription factor *Nanog*. Here, we aimed at identifying repressive histone modifications that would mark the *Nanog* locus for inactivation in response to ERK activity. We found histone H3 lysine 9 tri-methylation (H3K9me3) focally enriched between the *Nanog* promoter and its −5kb enhancer. While in undifferentiated ES cells H3K9me3 at *Nanog* depends on ERK activity, in somatic cells it becomes ERK-independent. Moreover, upon deletion of the region harbouring H3K9me3, ES cells display reduced heterogeneity of NANOG expression, delayed commitment into differentiation and impaired ability to acquire a primitive endoderm fate. We suggest that establishment of irreversible H3K9me3 at specific master regulators allows the acquisition of particular cell fates during differentiation.

## Introduction

Mouse Embryonic Stem (ES) cells are derived from preimplantation embryos and recapitulate numerous properties of the pluripotent inner cell mass of the blastocyst**^1^**. In vivo, the culmination of pluripotency – the ability to give rise to all three somatic germ layers – takes place when the primitive endoderm – a source of extra-embryonic tissues – segregates from the epiblast, the founder of the embryo proper**^2^**. This segregation is strictly controlled by the transcription factor *Nanog*, which is required to form the epiblast**^3,4^** and, additionally, stimulates FGF4 production. This extracellular signal is then transduced into neighbouring cells by ERK activity, silencing *Nanog* and opening a window of opportunity to undergo commitment into primitive endoderm differentiation**^5–10^**. This binary cell fate decision is characterised by substantial heterogeneity of NANOG expression, which creates the conditions required for epiblast and primitive endoderm specification**^10^**. Subsequently, *Nanog* expression is downregulated in the epiblast, eliciting the establishment of somatic differentiation**^11^**. In vitro, ES cells also exhibit extensive *Nanog* heterogeneity: while NANOG-positive cells self-renew efficiently, NANOG-negative cells are prone for differentiation even though they remain uncommitted and can spontaneously revert to the *Nanog*-expressing state**^12–16^**. Notably, NANOG-negative cells spontaneously generated in culture or by homozygous targeted deletion, show increased differentiation propensity towards both primitive endoderm and somatic fates**^11–16^**. NANOG heterogeneity has been proposed to result from several mechanisms. On the one hand, it has been suggested to result from the architecture and the topology of the pluripotency network**^17–19^**; on the other, it is known to strongly depend on extrinsic cues, such as LIF/STAT3, WNT/GSK3b and FGF/ERK signalling**^14,15,20,21^**. For both, specific regulatory properties and their inherent stochastic nature have been suggested to play a role**^22^**. Nevertheless, ERK activity has emerged as a key trigger of *Nanog* heterogeneity, in line with its general role in eliciting exit from pluripotency**^23,24^**. However, little is known about the chromatin-based aspects of *Nanog* heterogeneity. More specifically, it is unknown whether specific chromatin modifications**^25^** contribute to the stabilisation of the NANOG-negative state, which has been shown to be perpetuated for several cell divisions during self-renewal. Indeed, temporal analysis of NANOG fluctuations across cell generations has shown that the progeny of NANOG-negative cells is enriched in cells that maintain *Nanog* silencing, even though they can revert back and re-express NANOG**^26^**. This stability of the NANOG-negative state sets *Nanog* heterogeneity apart from other phenomena of gene expression heterogeneity, generally characterised by fast switching dynamics resulting from intrinsic and extrinsic noise or encoded in regulatory networks themselves**^27,28^**.

In this study, we aimed at identifying histone modifications associated with gene repression that would be: (i) differentially enriched at the *Nanog* locus when it is active or inactive; (ii) controlled by the signalling pathways associated with heterogeneity; (iii) heritable from mother to daughter cells**^29^**. We found histone H3 lysine 9 tri-methylation**^30^** (H3K9me3), enriched between the *Nanog* promoter and its-5kb enhancer**^31,32^**, to fulfil these criteria. Deletion of the region harbouring H3K9me3 at *Nanog* was associated with reduced NANOG heterogeneity, delayed commitment into differentiation and, most notably, defective differentiation along the primitive endoderm lineage. Moreover, our data also suggests that during differentiation, H3K9me3 at *Nanog* becomes independent from ERK activity. Hence, we propose that the timely establishment of ERK-independent H3K9me3 at *Nanog* marks commitment into differentiation and impacts cell fate acquisition in a lineage-dependent manner.

## Results

### ERK-dependent, mitotically stable, H3K9me3 at the *Nanog* locus in ES cells

To explore the involvement of chromatin marks potentially distinguishing active and inactive *Nanog* states, we first p erformed a c omparison of ES cell p opulations exhibiting heterogeneity, cultured in FCS+LIF **(Fig.1A)**, with highly homogeneous populations obtained by double inhibition of ERK and GSK3b (2i+LIF; **Fig.1A**). While several euchromatic marks associated with active transcriptional states were found more enriched in 2i+LIF, as expected, we observed a single repressive mark, H3K9me3, to be enriched in FCS+LIF and lost in 2i+LIF **(Fig.1B)**. H3K9me3 was present neither at the promoter (P in **Fig.1B**) nor at the enhancer (E in **Fig.1B**); rather, we detected H3K9me3 in the intervening region (IR in **Fig.1B**). To further characterise H3K9me3, we performed a high-resolution analysis of the *Nanog* locus (**Fig.1C**, top), which confirmed the presence of a robust peak of H3K9me3 between the enhancer and the promoter in FCS+LIF exclusively (**Fig.1C**). Analysis of total H3 confirmed the specificity of H3K9me3 enrichment (**Fig.1C**), which cannot be merely attributed to changes in nucleosome positioning or occupancy. Moreover, we observed good retention of H3K9me3 at *Nanog* in mitotic ES cells (**Fig.1C**), where drastic changes in nucleosomal densities could also be observed at the *Nanog* enhancer**^33^** (**Fig.1C**). Next, we used a previously described *Nanog*-GFP reporter**^11^** to sort *Nanog*-positive and -negative ES cells by FACS. We observed that H3K9me3 was more prominent in *Nanog*-negative cells, with clear spreading towards the promoter **(Fig.1D)**. Finally, we assessed the temporal ERK and GSK3b dependencies of H3K9me3 at *Nanog*. In 3 days of ERK inhibition with PD0325901, which induce high and homogeneous NANOG expression **(Fig.S1)**, H3K9me3 was significantly reduced (PD in **Fig.1E**), whereas even after 6 days of GSK3b inhibition with CHIR99021, H3K9me3 levels remained globally unchanged (CH in **Fig.1E**). Hence, the repressive H3K9me3 mark shows ideal properties to play a role in *Nanog* heterogeneity as it is readily dependent on ERK activity, fully lost in homogeneous NANOG populations, over-enriched in *Nanog*-negative cells, and maintained during mitosis.

**Fig. 1.**
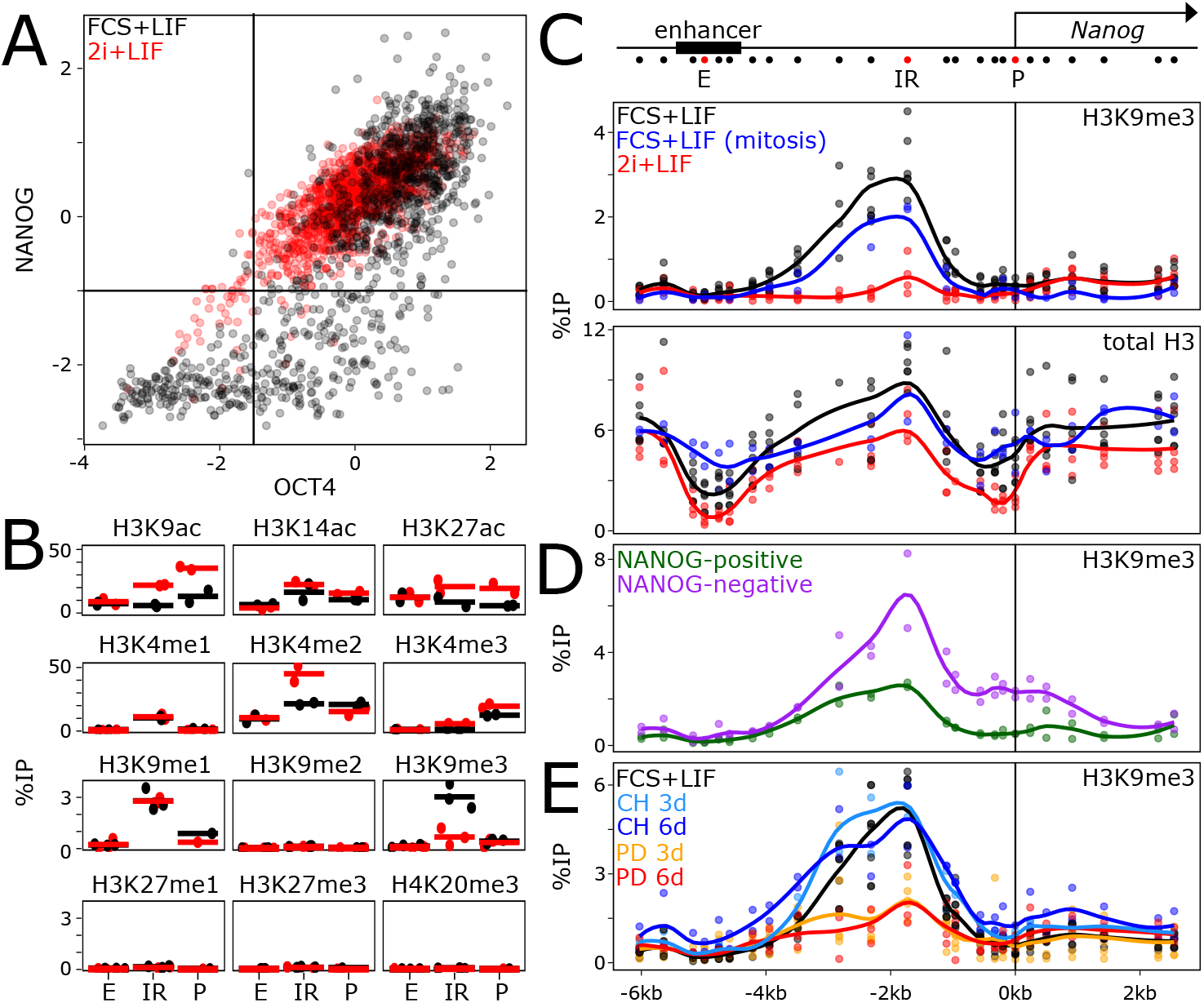
Mitotically stable, ERK-dependent H3K9me3 at the *Nanog* locus. **(A)** Quantification of OCT4 (z score; X-axis) and NANOG (z score; Y-axis) levels as assessed by immuno-staining in ES cells cultured in FCS+LIF (black, n=1125) and 2i+LIF (read, n=1445). **(B)** ChIPqPCR of several histone modifications as indicated, at three positions of the *Nanog* locus (the *Nanog* promoter, P; the *Nanog* −5kb enhancer, E; an intervening region, IR, located 1.7kb of the *Nanog* transcription start site; see map on (C)) in ES cells cultured in FCS+LIF (black) or in 2i+LIF (red). Each dot represents the percentage of immunoprecipitation (%IP; Y-axis) of independent replicates and the bar the corresponding mean. **(C-E)** Extended profile of H3K9me3 or total H3, as indicated, across the *Nanog* locus (X-axis represents genomic distances in kb with respect to the *Nanog* transcription start site, as schematised on top), in ES cells cultured as indicated in each plot. Each dot represents the percentage of immunoprecipitation (%IP; Y-axis) of independent replicates and the line the corresponding loess regression.

### H3K9me3 at *Nanog* impacts NANOG heterogeneity

To study the relevance of H3K9me3 at *Nanog*, we used a Crispr/Cas9 approach deleting 1.8kb between the enhancer and promoter (**Fig.2A**, red box). Two clones (ΔK9.1 and ΔK9.2) were confirmed as homozygously deleted with a complete absence of H3K9me3 **(Fig.2A)**. *Nanog* mRNA levels were slightly upregulated in ΔK9 cells **(Fig.2B)**, which presented a clear shift in NANOG expression to higher and more homogeneous levels, with a strong reduction of the proportion of cells expressing no or low NANOG **(Fig.2C)**. In agreement with the low upregulation of *Nanog* in ΔK9 cells, we observed a nearly insignificant increase in self-renewal efficiency, as determined by clonal assays. However, ΔK9 cells were more recalcitrant to efficiently differentiate upon LIF withdrawal **(Fig.2D)**. Next, we performed RNA-seq analysis to compare wild-type and ΔK9 cells **(Table S1)**, which confirmed a small increase in *Nanog* expression (FDR<0.05; **Fig.S2A**). We identified 235 and 402 genes that were up- or down-regulated, respectively, in both ΔK9 clones (FDR<0.05 and abs(log2FC)>0.3; **Fig.S2B**). Even though differentially expressed genes exhibited small fold changes **(Fig.S2C)**, consistent with the small increase in *Nanog* expression observed in ΔK9 cells, they were nonetheless found enriched in the vicinity of NANOG binding regions, compared with regions bound by OCT4/SOX2 but not NANOG**^33,34^ (Fig.S2D)**. We conclude that the deletion of the region harbouring H3K9me3 at the *Nanog* locus reduces the capacity of ES cells to explore the NANOG-negative state efficiently, leading to a measurable resistance to differentiate.

**Fig. 2.**
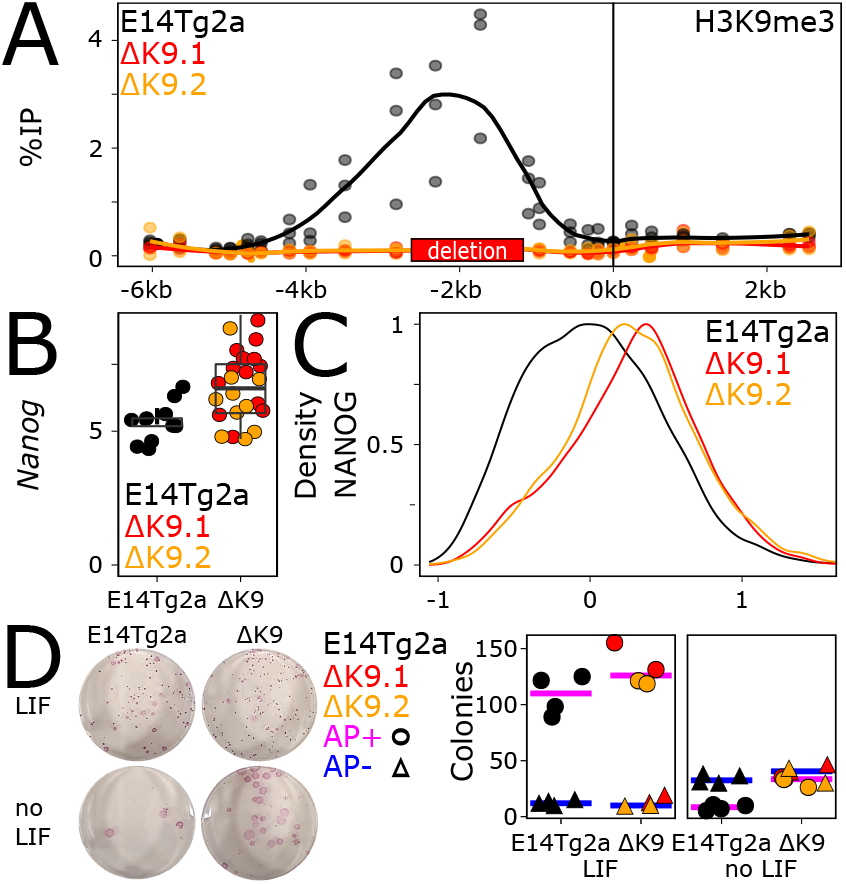
The *Nanog* region harbouring H3K9me3 enables heterogenous NANOG expression. **(A)** H3K9me3 profile across the *Nanog* locus, presented as in Figure 1C, in wild-type ES cells (E14Tg2a, black) and in two mutant derivatives (ΔK9.1 and ΔK9.2, in red and orange respectively), carrying a deletion of the region enriched for H3K9me3 (red box on the X-axis). **(B)** Expression of *Nanog* mRNA levels (normalised to *Tbp*) assessed by RT-qPCR in wild-type (E14Tg2a, black) and mutant clones (red and orange). Each dot represents the level of *Nanog* mRNA of independent replicates and the boxplots the corresponding median (bar), 25-75% percentiles (box) and 1.5-folds the inter-quartile range (whiskers). **(C)** Histogram representing the density (Y-axis) of NANOG expression levels (X-axis; log2(mean intensity) corrected to the E14Tg2a median of each experiment, n=2) in wild-type (E14Tg2a, black; n= 8191) or mutant cells (ΔK9.1 and ΔK9.2 in red and orange, respectively; n= 3835, 3812) as assessed by immuno-staining. **(D)** Representative alkaline-phosphatase staining of ES cell colonies for the indicated cell lines and culture conditions. The plot shows the number of alkaline-phosphatase (AP) positive (circles) and negative colonies (triangles) in wild-type (E14Tg2a, black) and ΔK9 cells (red and orange). Each dot represents an independent replicate and the bar the corresponding median for AP+ (pink) or AP-(blue) colonies.

### ERK-independent H3K9me3 at *Nanog* in somatic cells

We next aimed at characterising the status of H3K9me3 at *Nanog* in non-pluripotent cells. First, we established the H3K9me3 profiles over the *Nanog* locus in several cell types where *Nanog* has been silenced during development **(Fig.3A)**. H3K9me3 was systematically found enriched between the *Nanog* enhancer and promoter, albeit at different levels. In Trophectoderm Stem (TS) cells, the levels found at *Nanog* were lower than in FCS+LIF ES cells, except at the *Nanog* promoter where some spreading was detected. In Extra-embryonic Endoderm (XEN) cells, the levels of H3K9me3 were found higher, exhibiting spreading towards the promoter. Finally, in Mouse Embryonic Fibroblasts (MEFs), the level of H3K9me3 was particularly high, with prominent invasion of the *Nanog* promoter and gene body. Therefore, we conclude that while H3K9me3 is found at *Nanog* in the three cell types analysed, its absolute levels and the degree of spreading towards the promoter are variable. This suggests that a certain level of developmental specificity impacts H3K9 methylation at *Nanog*. Moreover, and in contrast to ES cells, the inhibition of ERK in MEFs did not abolish H3K9me3 at *Nanog*, which remained robustly enriched **(Fig.3A)**. This indicates that during differentiation, when *Nanog* silencing becomes irreversible, H3K9me3 at *Nanog* is liberated from its strict dependency on ERK.

**Fig. 3.**
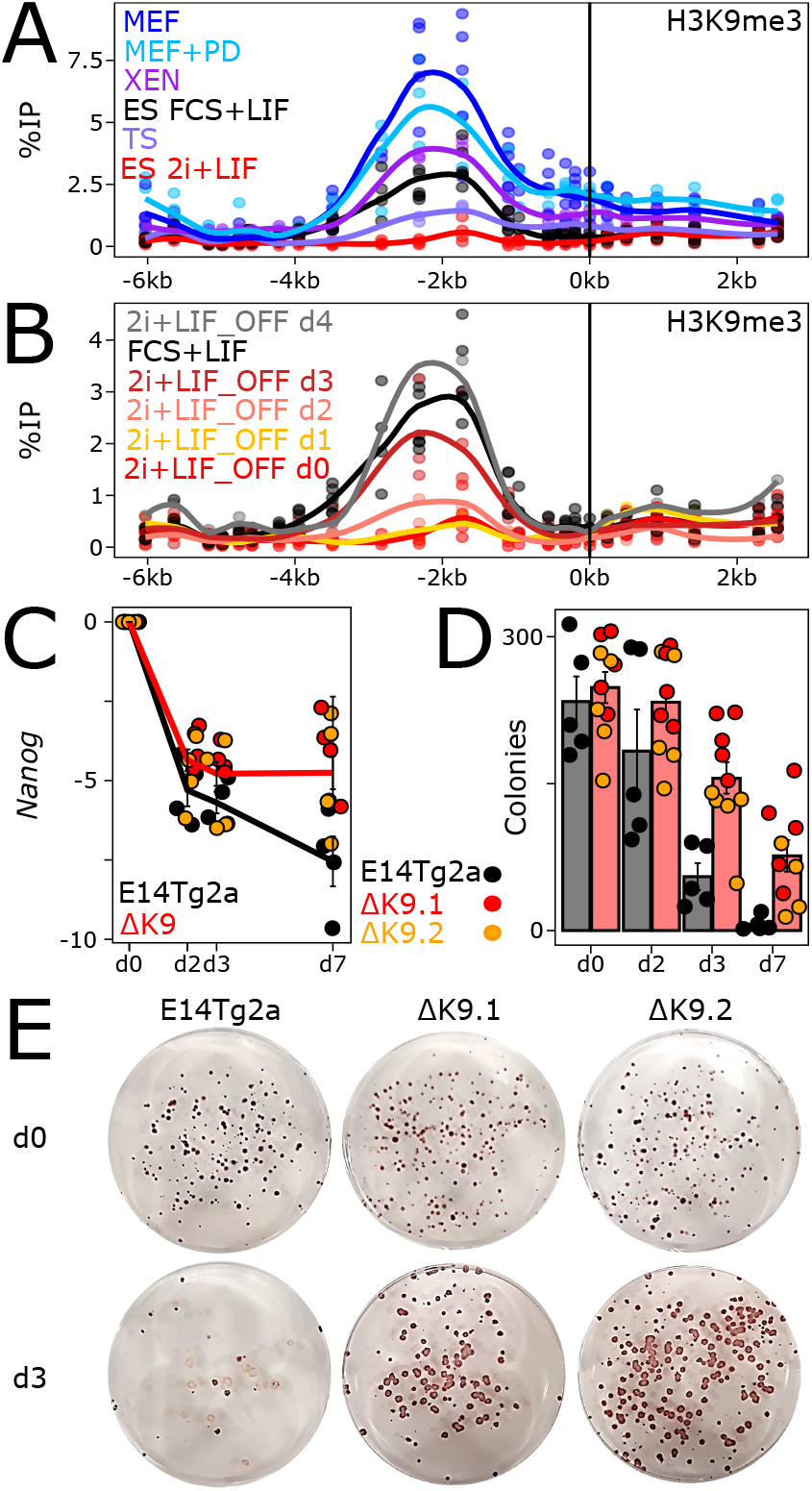
H3K9me3 at *Nanog* times commitment into differentiation. **(A)** H3K9me3 profile across the *Nanog* locus, presented as in Figure 1C, in the indicated cell lines and conditions. **(B)** Identical profiles for ES cells undergoing differentiation (labelled 2i+LIF_OFF) for the indicated number of days. **(C)** *Nanog* Log2 relative mRNA levels (d0 set to 1) measured by RT-qPCR and normalised to *Tbp*, during ES cell differentiation in the indicated cell lines. Each dot represents an independent replicate and the line the corresponding mean with standard errors. **(D)** Number of alkaline-phosphatase positive colonies obtained after switching to 2i+LIF wild-type (E14Tg2a, black) and ΔK9 (red and orange) cells seeded clonally and differentiated for the number of days indicated on the X-axis. Each dot represents an independent replicate and the histogram the corresponding mean and standard error. **(E)** Representative alkaline-phosphatase staining of ES cell colonies cultured in 2i+LIF after 0 (top) or 3 (bottom) days of differentiation for the indicated cell lines.

### H3K9me3 at *Nanog* controls the timing of commitment into differentiation

We then aimed at assessing the dynamics of H3K9me3 at *Nanog* during ES cell differentiation. We used a simple protocol starting from 2i+LIF (absence of H3K9me3) and based on the withdrawal of LIF and of ERK and GSK3b inhibitors **(Fig.S3A)**. We observed a step-wise increase of H3K9me3 **(Fig.3B)**: if it remained low during the first 48h, it suddenly appeared after 3 days and only slightly increased after 4 days. Somehow unexpectedly, this appearance of H3K9me3 did not correlate with a particularly strong reduction of *Nanog* expression **(Fig.3C)**. In fact, we observed *Nanog* downregulation taking largely place during the first 48h, in the absence of high levels of H3K9me3. However, while *Nanog* expression continued to decrease during differentiation of wild-type cells, we observed that ΔK9 clones displayed a stabilisation of low *Nanog* expression after the sharp decrease occurring during the first 2 days **(Fig.3C)**, despite an efficient differentiation **(Fig.S3A)**. This different behaviour of *Nanog* expression in ΔK9 clones, temporally correlated with the time at which H3K9me3 is established in wild-type cells, prompted us to determine if commitment into differentiation – the moment at which cells cannot easily come back to an undifferentiated state – was altered in ΔK9 cells. For this, we seeded wild-type and ΔK9 cells at clonal density and after 2, 3 or 7 days of differentiation, we replaced the culture medium back to 2i+LIF: only cells that have not yet irreversibly lost their capacity to self-renew will survive, proliferate and form undifferentiated, alkaline-phosphatase-positive colonies**^35^ (Fig.3D,E)**. In wild-type cells, we observed a striking coincidence of the time of commitment, taking place between days 2 and 3, with the appearance of H3K9me3 at *Nanog*. In ΔK9 clones, however, we observed a significant number of alkaline-phosphatase-positive colonies after 3 and even 7 days of differentiation, testifying of delay in commitment. Altogether, these analyses suggest that in 2i+LIF ES cells, H3K9me3 at *Nanog* is established during differentiation, when it marks the irreversible commitment into effective differentiation. Yet, it is not strictly required for differentiation per se and rather enables the appropriate timing of commitment.

### ΔK9 cells exhibit delayed differentiation

Given the delay in differentiation commitment observed in ΔK9 clones, we monitored the expression of several genes reflecting the loss of pluripotency and the transition to a differentiated state**^35^ (Fig.S3B)**. While naïve pluripotency genes (*Esrrb, Klf4, Prdm14, Rex1*) showed a less drastic downregulation, mimicking *Nanog* expression, differentiation markers (*Fgf5, Dnmt3b, Otx2, Wnt3*) showed delayed dynamics. Next, we differentiated wild-type and ΔK9 cells into Embryoid Bodies (EBs), a paradigm that recapitulates the establishment of multiple lineages. At the morphological level we observed ΔK9 EBs to be often characterised by defective sealing at their periphery **(Fig.S4A)**. Moreover, cellular outgrowths derived from ΔK9 EBs also exhibited obvious differences compared to those derived from wild-type EBs, with less apparent multi-lineage differentiation **(Fig.S4A)**. Gene expression analyses of EBs after d4, d6 and d8 of differentiation confirmed the attenuated downregulation of *Nanog* expression **(Fig.4A)**. Moreover, PCA analysis of RNA-seq profiling **(Table S1)** highlighted a transcriptome-wide delay of both ΔK9 clones, starting at d4 and progressively increasing through time **(Fig.4B)**. Gene ontology analysis of the top 1000 loadings of the PCA identified focal adhesion genes among the most enriched cellular components (GO:0005925; p<10**^−5^**), in line with our morphological observations **(Fig.S4A)**. Therefore, the lack of H3K9me3 at *Nanog* is strongly associated with delayed differentiation, as evaluated with three distinct differentiation paradigms **(Fig.2D, Fig.S3B and Fig.4B)**.

**Fig. 4.**
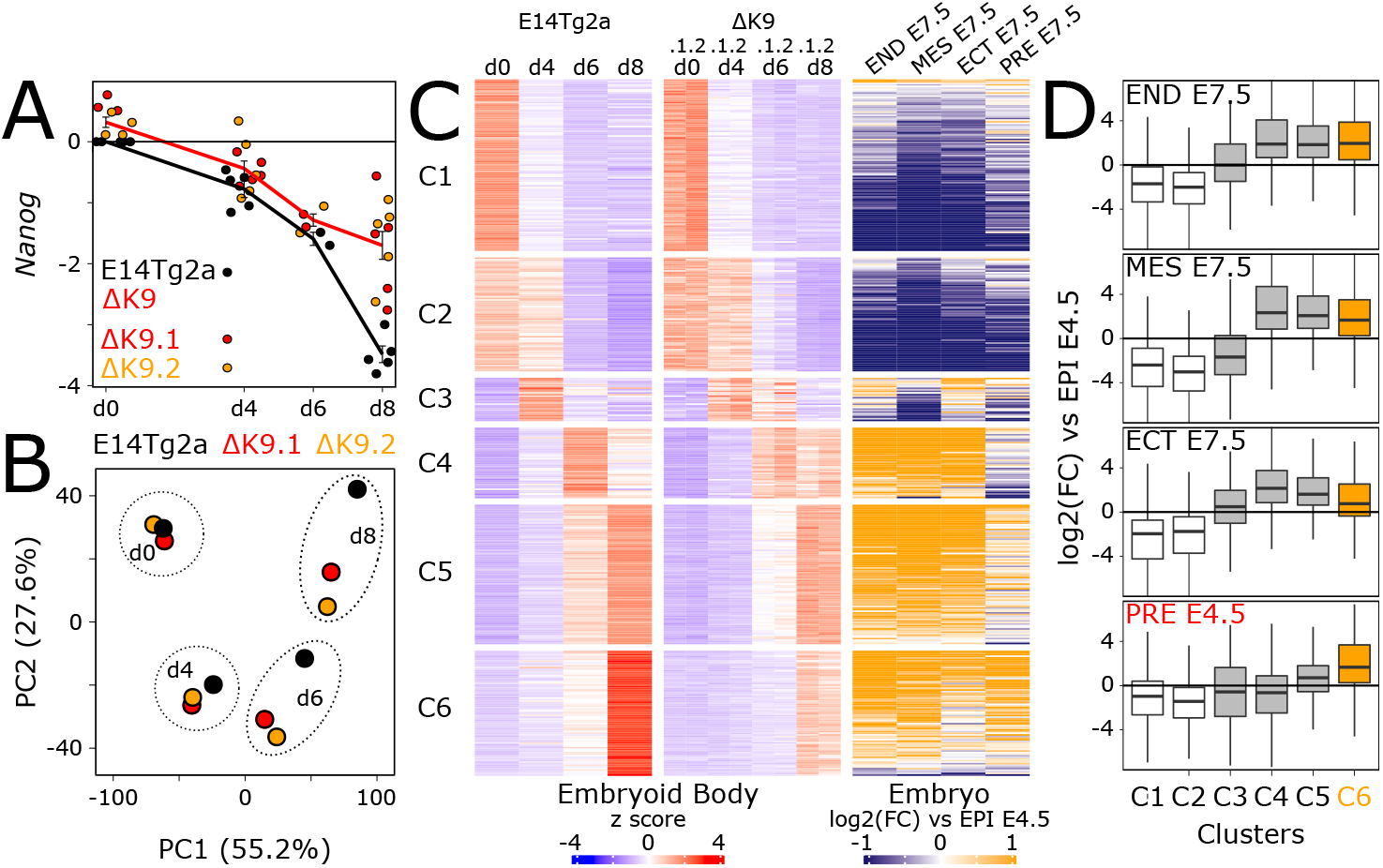
The lack of H3K9me3 at *Nanog* leads to delayed differentiation. **(A)** Log2 *Nanog* mRNA levels measured by RT-qPCR and normalised to *Tbp* during embryoid body differentiation in the indicated cell lines. Each dot represents an independent replicate and the line the corresponding mean with standard errors. **(B)** Principal Component Analysis of 16336 transcripts quantified by RNA-seq (Transcripts Per Million above 1 in at least one sample) in wild-type (E14Tg2a, black; n=3) and ΔK9 (red and orange; n=3 for each) cells undergoing embryoid body differentiation for the indicated number of days. **(C)** Heatmap of 4100 transcripts displaying significant gene expression changes during embryoid body differentiation (DEseq FDR<0.05 and abs(log2FC)>1 in at least one comparison to undifferentiated cells; n=3 for each cell line), clustered by k-means (k=6; 50 iterations; 50 random starts) after z score normalisation. Left, relative gene expression (z score) in wild-type (E14Tg2a) and ΔK9 cells during embryoid body differentiation; right, corresponding log2 fold-changes of each developmental stage indicated on top versus E4.5 epiblast, as previously reported**^36^**. END: endoderm; MES: mesoderm; ECT: ectoderm; PRE: primitive endoderm; EPI: epiblast. **(D)** Boxplot (median; 25-75% percentiles; 1.5-folds the inter-quartile range) of log2FC shown in (C) for each cluster. Cluster C6 is highlighted in orange.

### The absence of H3K9me3 at *Nanog* leads to major defects in primitive endoderm differentiation

To further explore the molecular nature of the defects observed during EB differentiation, we first identified around 4000 genes displaying temporal gene expression changes in either differentiating wild-type or mutant cells compared to their undifferentiated controls (FDR<0.05, abs(log2FC)>1; **Table S1**). These genes were then clustered using k-means; this allowed us to identify 6 groups of genes displaying different expression dynamics during EB differentiation (**Fig.4C**, left; **Table S1**). While all identified clusters underscored the delay into differentiation, one in particular, cluster c6, showed a blatant attenuation of gene upregulation at d8. To characterise each cluster relative to known developmental trajectories, we plotted the fold-change reported in a previous study**^36^** where embryonic endoderm, mesoderm and ectoderm cells of E7.5 embryos, as well as primitive endoderm cells of E4.5 embryos, were directly compared to the E4.5 epiblast (**Fig.4C**, right). While clusters c1 and c2 showed a marked downregulation upon epiblast differentiation into all lineages, cluster c3 to c6 displayed a clear upregulation in at least one E7.5 lineage. Notably, cluster c6, which shows the strongest ΔK9 vs wild-type differences, was the only one enriched for genes displaying a prominent upregulation in the primitive endoderm with respect to the epiblast **(Fig.4D)**. Analysis of primitive endoderm markers such as *Gata6*, *Gata4* and *Sox17* during EB differentiation confirmed the altered induction of these genes **(Fig.S4B)**.

Finally, in light of these results, we wanted to ascertain whether the defective primitive endoderm signature identified in EBs implies a deficiency in the capacity of ΔK9 clones to engage in primitive endoderm differentiation. Thus, we challenged wild-type and ΔK9 ES cells with a primitive endoderm differentiation protocol**^37^ (Fig.S5)**. In wild-type cells, but not in ΔK9 clones, we observed the appearance of endoderm-like cell clusters from day 4 onwards. Moreover, ΔK9 clones showed increased cell death. Immunofluorescence of NANOG, GATA6, GATA4 and PDGFRa, confirmed NANOG silencing in cells expressing primitive endoderm markers, as expected **(Fig.5A,B,C)**. In ΔK9 clones, however, NANOG expression was prominent and the appearance of cells expressing primitive endoderm markers anecdotical. Therefore, ΔK9 cells are not able to undergo directed differentiation into primitive endoderm. We conclude that H3K9me3 is required to stably silence *Nanog* during differentiation and that failing to do so has different consequences depending on the differentiation lineage, with primitive endoderm being particularly sensitive.

**Fig. 5.**
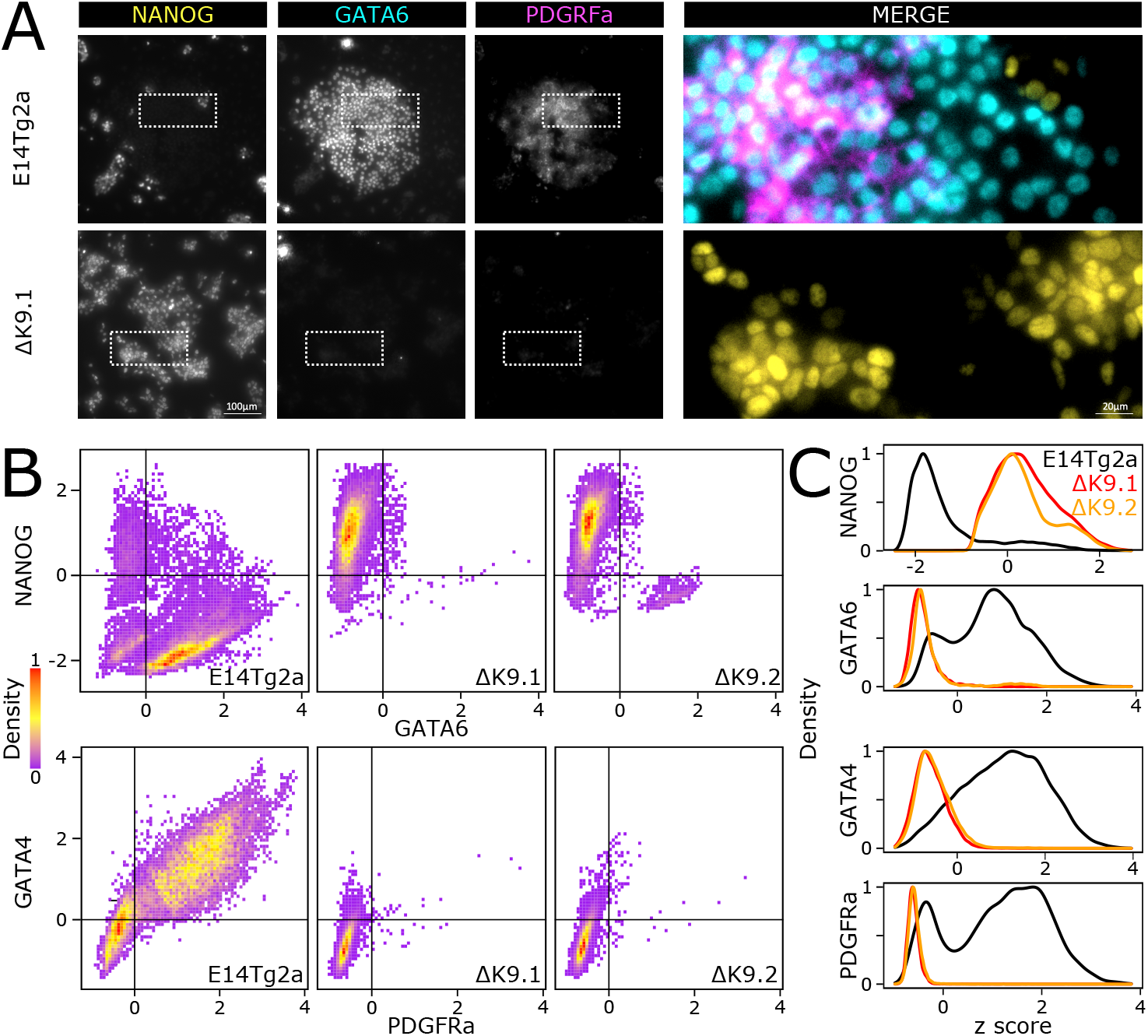
The lack of H3K9me3 at *Nanog* abolishes primitive endoderm differentiation. **(A)** Representative immuno-staining of ES cells (E14Tg2a, top; ΔK9.1, bottom) subject to a directed differentiation protocol into primitive endoderm. **(B-C)** Quantification of dual immuno-staining for NANOG, GATA6 and GATA4, PDGRFa in E14Tg2a (n=19507, 20641), ΔK9.1 (n=11348, 19476) and ΔK9.2 (n=9713, 19742) cells differentiated as in (A).

## Discussion

Gene expression heterogeneity has emerged as a main motor of lineage diversification during development, particularly in stem cell and progenitor populations: upon appropriate stimuli, these expression differences are translated into an effective decision-making process that culminates in commitment**^22,27,28^**. When these heterogeneities are dynamic, such as in the case of *Nanog* and ES cells, the fluctuating expression is eventually fixed. A likely model accounting for the transition from reversible to irreversible silencing involves epigenetic repression, in the sense of heritable chromatin states incompatible with transcription that do not depend on the original triggers**^38^**. The acquisition of epigenetic repression, including H3K9me3**^39^**, is indeed perceived as a general mean to restrict developmental fate choices as cells differentiate**^40^**. Accordingly, we show that *Nanog* silencing during differentiation is accompanied by H3K9me3, which irreversibly locks its expression and thus contributes to commitment into differentiation. Two observations, however, indicate that additional complexity may characterise both *Nanog* silencing and its developmental implications.

While epigenetic repression is often instated de novo during differentiation, for example at *Oct4***^41^**, the regulation of *Nanog* appears to involve an intermediary state where H3K9me3 is already established but not yet fixed. In undifferentiated cells, H3K9me3 is readily detected at *Nanog*, particularly in *Nanog*-negative cells. Moreover, it is transferred from mother to daughter cells during mitosis. However, it is strictly dependent upon ERK signalling, the main driver of NANOG heterogeneity. This mitotically-stable and ERK-dependent state of H3K9me3 confers to *Nanog* silencing the required stability to be inherited and, at the same time, sufficient flexibility to revert back to transcriptional activity. During differentiation (at least as judged by the analysis of embryonic fibroblasts), H3K9me3b ecomes independent of ERK and, with respect to ERK, irreversible. Therefore, even if subjected to variations in ERK stimuli, *Nanog* will remain silent. In spite of these considerations, we suggest that H3K9me3 at *Nanog* becomes epigenetic exclusively during differentiation, when it is not any longer dependent on its original trigger. Whether this transition is mediated by direct mechanisms operating at the locus, on the H3K9me3-associated machinery, or linked to the general lack of strong epigenetic repression in ES cells**^42,43^**, remains unknown. Nevertheless, these findings are reminiscent of the dependency of other repressive marks, such as H3K27me3, on LIF signalling and NANOG activity**^34^**, suggesting a general vassalage of repressive chromatin marks to more dynamic regulators in undifferentiated ES cells. By displaying regulated dependencies towards signalling and/or transcription factor activity, repressive chromatin modifications may facilitate conditional heritability and excitability or, on the contrary, fixed gene expression states**^42^**.

*Nanog* is known to counteract differentiation when ectopically expressed at high levels**^11^**. Since the deletion of the region harbouring H3K9me3 leads to a minor increase of NANOG expression, it was not expected to block differentiation. After all, upon the collapse of the pluripotency network triggered by differentiation signals, *Nanog* would lose most of its activators and be downregulated, as we observed. Yet, the lack of H3K9me3 leads to a lack of complete *Nanog* silencing, providing an opportunity to evaluate the importance of fully repressing *Nanog* during differentiation. Similarly, as cells lacking H3K9me3 at *Nanog* reduce their heterogeneity in a context where *Nanog* can nevertheless be downregulated (in contrast to ectopic expression systems), the importance of NANOG heterogeneity in lineage priming can also be inferred from our experimental setup. We observed delayed commitment and altered differentiation into all germ layers. Yet, the highest consequences affect genes normally upregulated in the primitive endoderm; remarkably, the lack of H3K9me3 at *Nanog* is incompatible with differentiation along this lineage. This observation underscores the relative importance of *Nanog* in repressing differentiation of somatic versus primitive endoderm lineages in ES cells. Furthermore, *Nanog* heterogeneity in ES cells has been proposed to either reflect the heterogeneity observed in early blastocysts, which cells of the inner mass can either express NANOG or GATA6, or the early downregulation of *Nanog* taking place around implantation to elicit somatic differentiation events of the epiblast**^11–16^**. Indirectly, thus, our results could be interpreted as *Nanog* heterogeneity and its subsequent full silencing being functionally associated to the epiblast versus primitive endoderm specification. In this regard, two observations are noteworthy. First, the repressive H3K27me3 mark has been shown to play a preponderant role in downregulating genes that prime ES cells for primitive endoderm differentiation**^44^**. Second, *Nanog* sustains H3K27me3 in undifferentiated and early differentiating ES cells**^34^**. In light of these findings and of our observation that the loss of H3K9me3 also alters the differentiation balance between somatic and primitive endoderm lineages, we suggest that a signalling and transcription factor dialogue established through repressive histone methylation contributes to the epiblast versus primitive endoderm specification. In this model, ERK dynamically controls H3K9me3 at *Nanog*, which sustains H3K27me3 levels and keeps primitive endoderm genes in check, generating reversible and mosaic expression patterns associated with either epiblast or primitive endoderm fates.

Overall, our observations add to the notion thatheterochromatin contributes to cell fate restriction during differentiation processes. They also suggest that these events are more nuanced in their action than it was anticipated, given the regulation of its dependency towards signalling cues in respect of its epigenetic potential and the differential impact that the ensuing stability of the repression may have in different lineages. Whether our findings can be extrapolated to other master regulators of pluripotency or to other developmental transitions represent important new avenues for the future.

## Supplementary information

Five supplementary figures accompany this manuscript, they can be found at the end of this document. Two Supplementary Tables and Methods are available online.

## Acknowledgements

This study was conceived by P.N with inputs from A.D. Experiments were designed and executed by A.D., with help from L.V., S.V-P. and M.C-T. Computational analyses were done by A.C. and P.N. The paper was written by P.N. with inputs from A.D. and M.C.T. The authors acknowledge the cytometry platform of Institut Pasteur for technical assistance and L. Bally-Cuif and S. Tajbakhsh for critical reading of the manuscript. P.N. and M.C-T. acknowledge the Labex Revive (Investissement d’Avenir; ANR-10-LABX-73), the Institut Pasteur and the CNRS for funding.

---

**Supplementary Information, Fig. S 1.**
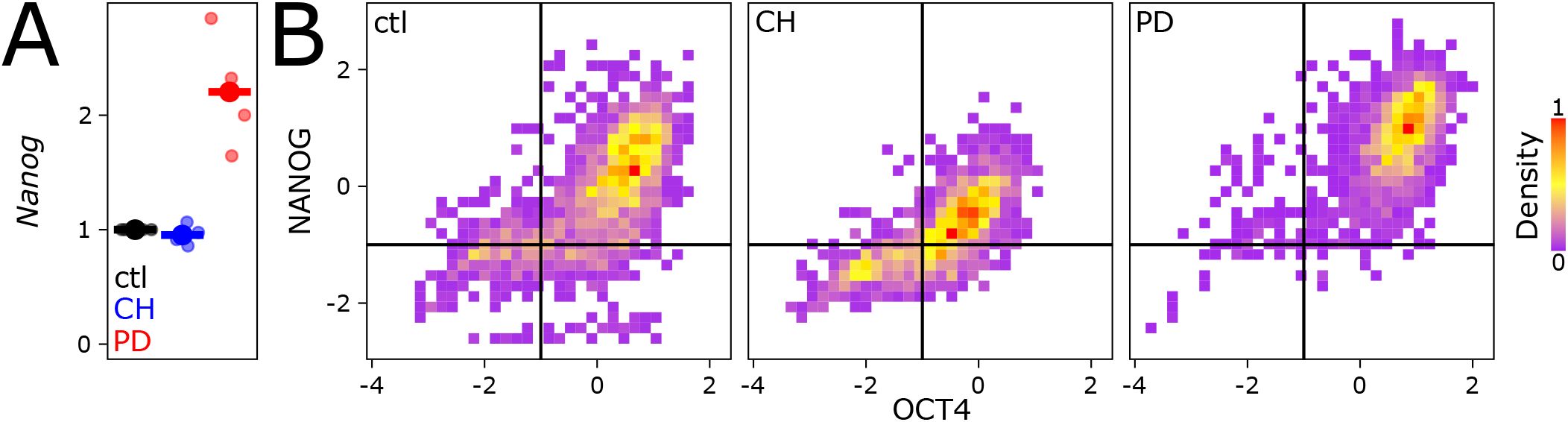
Effects of PD and CH treatment in ES cells. **(A)** Relative mRNA levels (ctl set to 1) of *Nanog*, measured by RT-qPCR and normalised to *Tbp*, upon 3d of PD or CH treatment of FCS+LIF cultured cells. **(B)** Quantification (z score) of NANOG and OCT4 immuno-staining in untreated E14Tg2a cells (ctl; n=2053) and after 3d of CH (n=1554) or PD (n=1857) treatment, as indicated.

**Supplementary Information, Fig. S 2.**
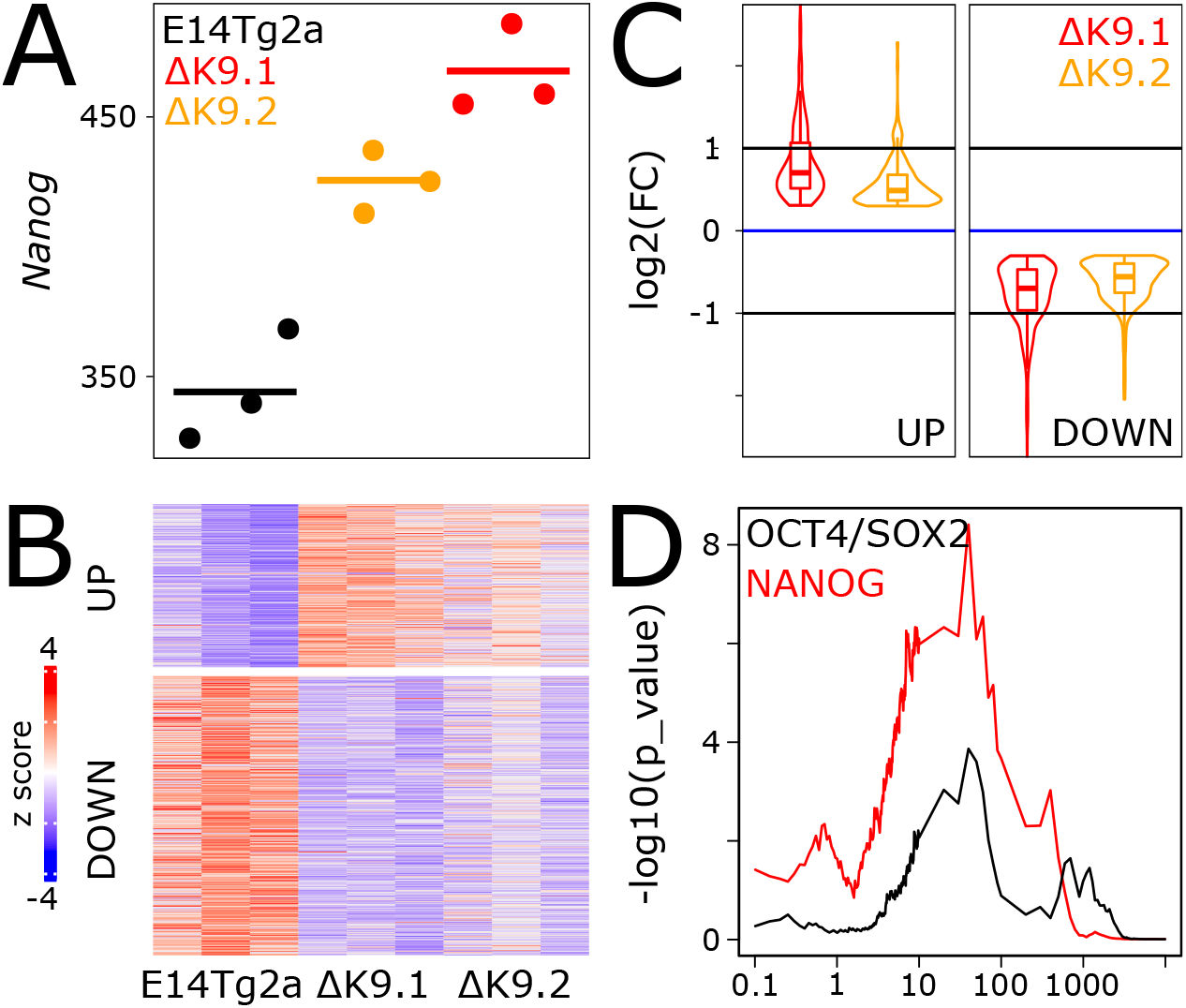
Gene expression consequences in ΔK9 cells. **(A)** Confirmation of *Nanog* upregulation in ΔK9 cells by RNA-seq (Transcripts Per Million, TPM). **(B)** Z scored heatmap of genes identified as differentially expressed in ΔK9 cells. **(C)** Violin-Boxplot of gene expression fold-changes measured in ΔK9.1 and ΔK9.2 clones compared to wild-type E14Tg2a cells. **(D)** Statistical association (Y-axis, -log10(Fisher exact test p-value)) between the differentially expressed genes shown in (B) with the presence of NANOG (red) or OCT4/SOX2 (black) binding sites as identified by ChIP-seq in a previous study**^33^**, as a function of the distance (X-axis, kb).

**Supplementary Information, Fig. S 3.**
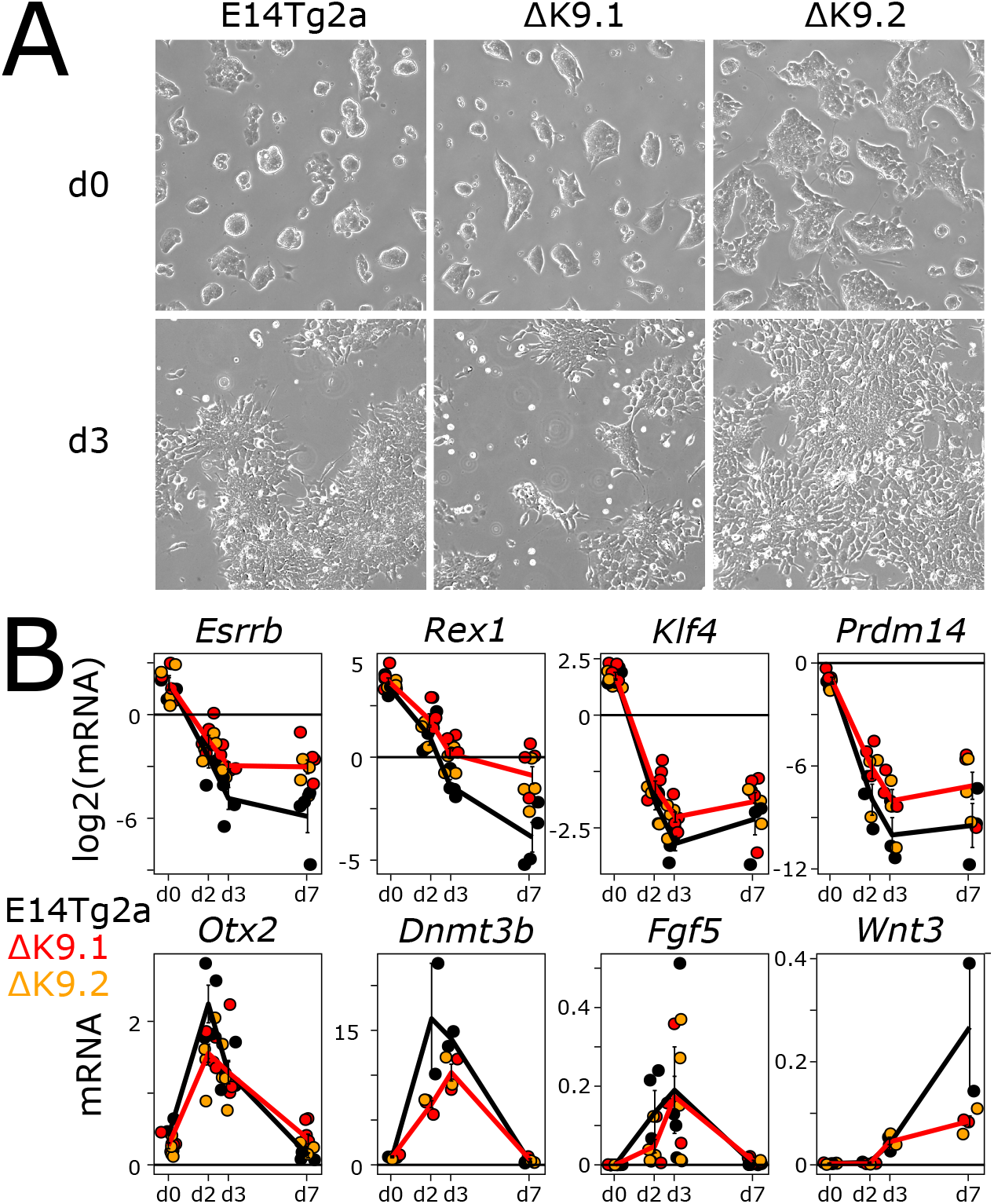
Differentiation of wild-type and ΔK9 cells upon 2i+LIF withdrawal. **(A)** Representative bright-field photomicrographs of wild-type (E14Tg2a) and ΔK9 cells cultured in 2i+LIF (top) and after 3 days of withdrawal (bottom). **(B)** RT-qPCR analysis of markers of naïve pluripotency (top) and differentiation (bottom) after 0, 2, 3 and 7 days of 2i+LIF withdrawal. *Tbp* was used for normalisation.

**Supplementary Information, Fig S 4.**
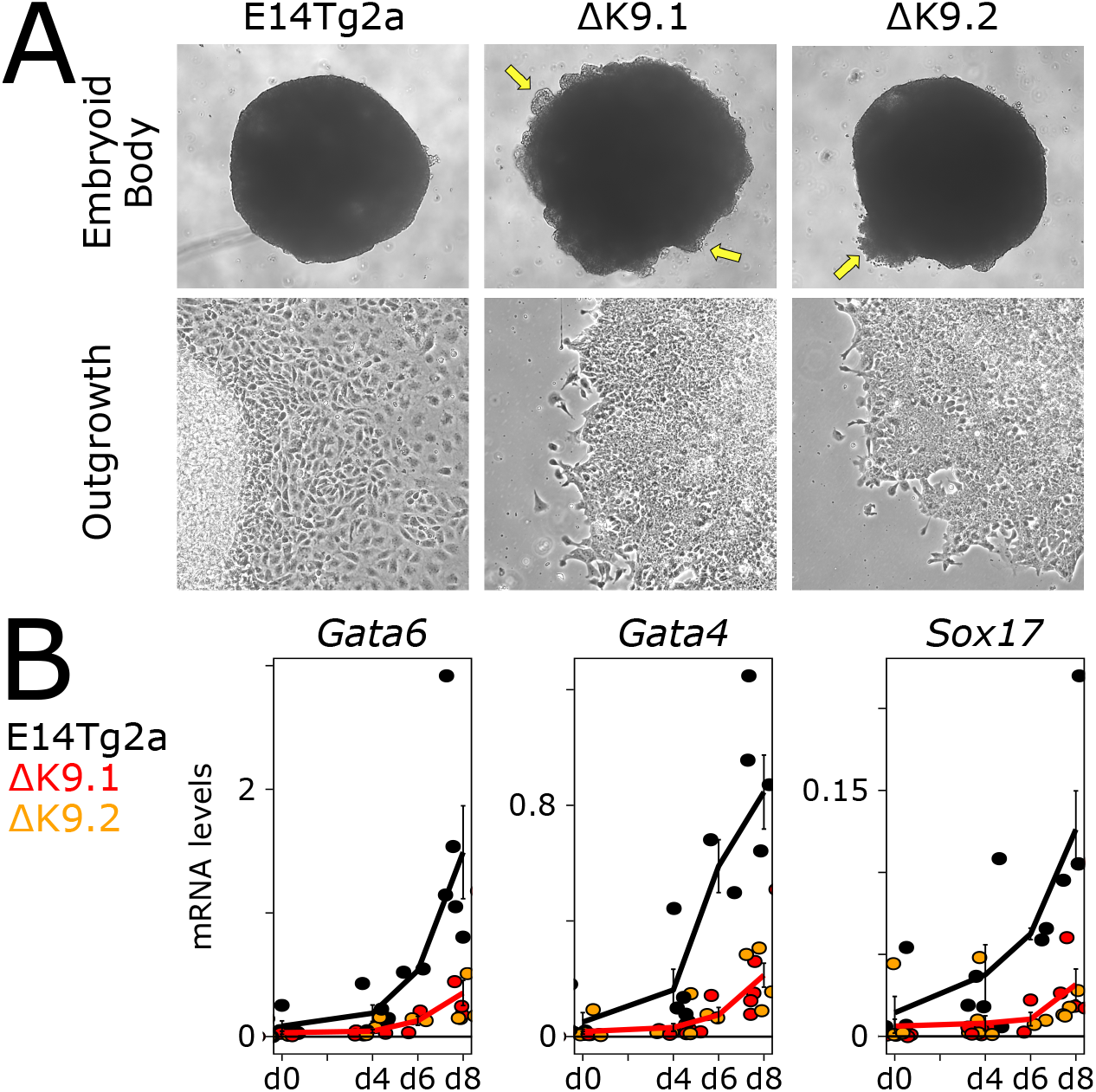
Differentiation of wild-type and ΔK9 cells into embryoid bodies. **(A)** Representative bright-field photomicrographs of wild-type (E14Tg2a) and ΔK9 embryoid bodies (top) and their derived cellular outgrowths (bottom). The arrows point to defective sealing of EBs periphery. **(B)** RT-qPCR analysis of endoderm markers after 0, 4, 6, 8 days of embryoid body differentiation. *Tbp* was used for normalisation.

**Supplementary Information Fig S 5.**
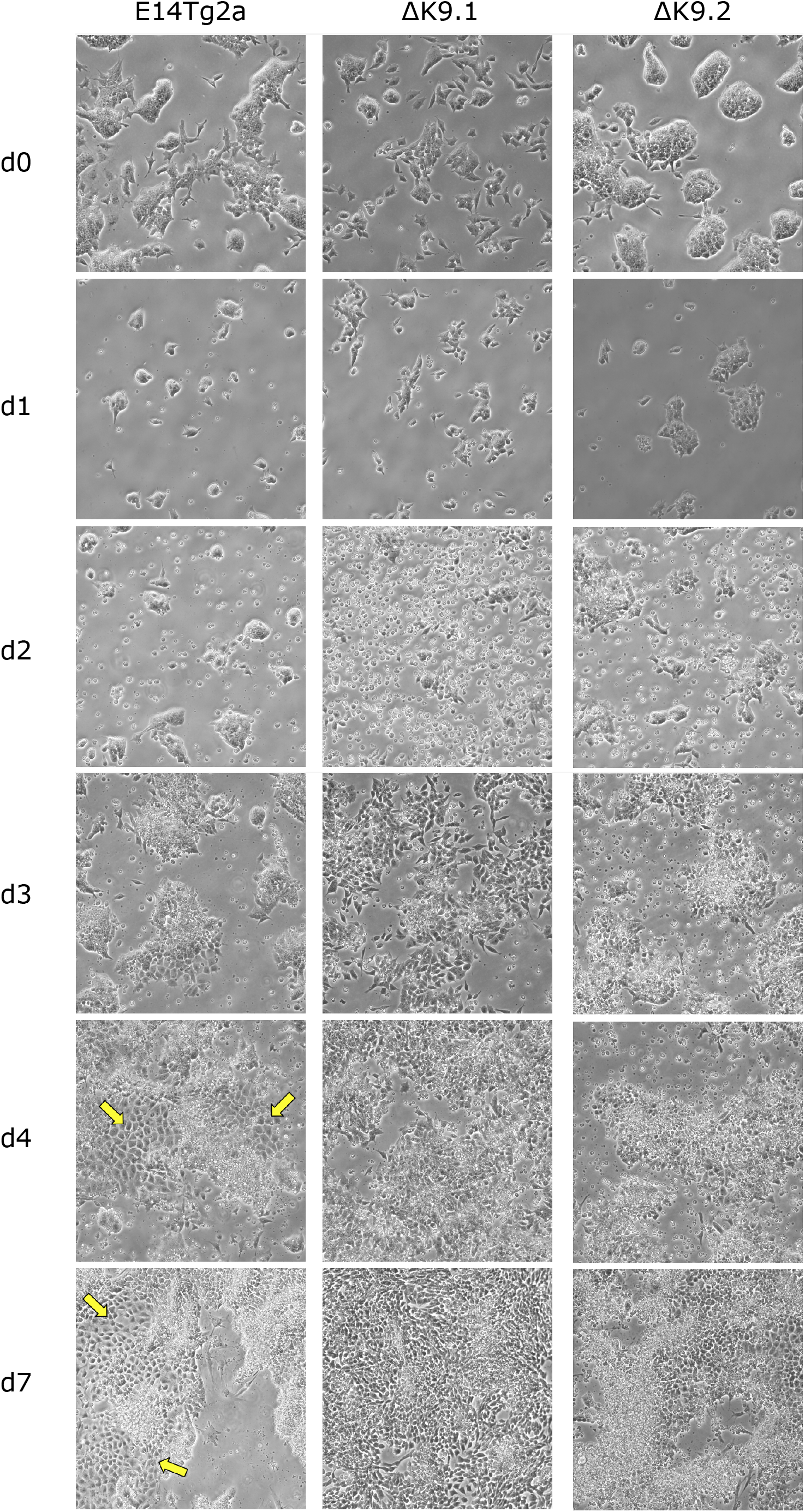
Directed differentiation of wild-type and ΔK9 cells into primitive endoderm. Representative bright-field photomicrographs of wild-type (E14Tg2a) and ΔK9 cells during their treatment with a primitive endoderm differentiation protocol. The arrows point to clusters of primitive endoderm cells.

