## Supplementary material for "H3K9 tri-methylation at *Nanog* times differentiation commitment and enables the acquisition of primitive endoderm fate": Methods

**Dubois et al.**

Contents:


1/ Cell culture page 2

2/ Flow cytometry page 5

3/ Imaging page 5

4/ Chromatin immunoprecipitation page 7

5/ Gene expression page 8

6/ References page 10


***

**1/ Cell culture.**

*1a/ General ES cell culture conditions.*

ES cells (E14Tg2a, ΔK9 clones and TNG, a kind gift of Pr. Ian Chambers) were cultured (37°C, 7%CO2) on 0.1% gelatine (SIGMA, Cat# G1890-100G) in DMEM, high glucose, GlutaMAX™ Supplement, pyruvate (Gibco, Cat# 31966-021), 10% FCS (Sigma, F7524), 100 µM 2-mercaptoethanol (Gibco, Cat# 31350-010), 1× MEM non-essential amino acids (Gibco, Cat# 1140-035) and 10 ng/ml recombinant LIF (MILTENYI BIOTEC, Cat# 130-099-895). When indicated, cells were grown in serum-free 2i-containing medium (1µM PD0325901 and 3 µM CHIR99021; Axon1408 & Axon1386 respectively): 0.5× DMEM/F12 (Gibco, Cat# 31331093), 0.5×Neurobasal (Gibco, Cat# 21103049), 0.5X N2 supplement 100X (Gibco, Cat# 17502048), 0.5X B27 supplement 50X (Gibco, Cat# 17504044), 10µg/ml Insulin (Sigma, Cat# I1882-100MG), 2 mM L-Glutamine (Invitrogen, Cat# 91139), 37.5 µg/ml BSA (Sigma, Cat# A3311-10G), 100 µM 2-mercaptoethanol (Gibco, Cat# 31350-010), 10 ng/ml recombinant LIF (MILTENYI BIOTEC, Cat# 130-099-895). Cells were seeded at 0.02M to 0.04M cells/$cm^2$, media changed every 2 days and cells passaged every 3-4 days. All 2i+LIF analyses were performed after at least 3-4 passages in 2i+LIF. All experiments inhibiting ERK or GSK3b were performed with 1µM PD0325901 and 3 µM CHIR99021, respectively. Cells were karyotyped and regularly tested mycoplasma-free.

*1b/ Culture conditions of MEFs, XEN and TS cells.*

Mouse Embryonic Fibroblasts (MEFs) were derived from F1 129sv/129sv E13.5 male embryos and cultured (37°C, 7%CO2), for no more than 4 passages, in DMEM, high glucose, GlutaMAX™ Supplement, pyruvate (Gibco, Cat# 31966-021), 10% FCS (Sigma, F7524), 100 µM 2-mercaptoethanol (Gibco, Cat# 31350-010), 1× MEM non-essential amino acids (Gibco, Cat# 1140-035). When indicated, MEFs were treated for 3 days with 1 µM PD0325901. Extra-Embryonic Endoderm (XEN) cell lines (XEN_ 7.9, XEN_ 7.3 and XEN_ 7.7) were initially isolated from Tgn ($X^{GFP}$) 4 Nagy ×129$Hprt^{bm1}$Pgk1a blastocysts (Kunath et al., 2005) and were routinely cultured (37°C, 7%CO2) in DMEM, high glucose, GlutaMAX™ Supplement, pyruvate (Gibco, Cat# 31966-021), 10% FCS (Sigma, F7524), 100 µM 2-mercaptoethanol (Gibco, Cat# 31350-010), 1× MEM non-essential amino acids (Gibco, Cat# 1140-035) on 0.1% gelatine, without LIF, and passaged every 4 days. Trophectoderm Stem (TS) cell lines (F2 and F3) were initially isolated from F1 129/Sv Hprt-4 Pgk1a ×129/Sv embryos (Kunath et al., 2005) and cultured (37°C, 5%CO2) in RPMI 1640 Medium, GlutaMAX™ Supplement (Gibco, Cat# 61870036) with 20% FCS (Sigma, F7524), 1 mM sodium pyruvate, 100µM β-mercaptoethanol (Gibco, Cat# 31350-010),

25 ng/ml hrFGF4 (235-F4-025, R&D) and 1 µg/mL heparin (H3149-10KU, Sigma) on mitomycined-MEFs. Cells were passaged every 2-3 days. For TS analyses, MEFs were removed by adsorption onto gelatinized culture dishes for 1.5 hours.

*1c/ Derivation of ΔK9 cells.*

gRNAs (5'- CAGAGGAGGGCTTAAGAGAT and 5'- CACTCTAACCCAGCTTAAGT) were designed and cloned under the control of a U6 promoter in a vector conferring puromocycin resistance, as described (Heurtier et al., 2019). 1µg of each gRNA-expressing vector and of a Cas9/mCherry expression vector (Addgene#64324) were lipofected in E14Tg2a cells according to manufacturer's instructions (Lipofectamine 2000; ThermoFisher). Lipofected cells were selected with Puromycin (1 µg/ml) and FACS-sorted for mCherry fluorescence. Puromycin-resistant and mCherry-positive cells were seeded at clonal density and ~100 clones were picked 10 days later. Clones were screened by 2 independent PCR (LongAmp Taq PCR kit; BioLabs Cat# E5200S) with primers spanning the deletion (DelCTRL_F/R; N12c_F/N17b_R; Table S2), by real-time PCR using primers along the *Nanog* locus (Table S2), and by cloning and sequencing of PCR products (N12c/N17b). Two karyotypically normal independent clones, ΔK9.1 and ΔK9.2, were selected for this study.

*1d/ Clonal assays.*

600 cells were plated in single wells of 6-well plates coated overnight with poly-L-ornithine 0.01% (Sigma, Cat# P4957) at 37°C, washed twice with PBS 1X and coated 2 h with 1X laminin (Sigma, Cat# L2020). After 7 days, cells were fixed and stained using an alkaline phosphatase staining kit (Sigma Aldrich, Cat # 86R-1KT) according to the manufacturer's instructions or by fixation 20 min in PFA 4%, 2 washes with PBS 1X, incubation 15 min at RT in the dark in a staining solution (TrisMaleate 1M, MgCl2 1M, 100 mg/ml $\alpha$-naphtyl-phosphate, 100 mg/ml Fast-Red TR), 2 washes with PBS 1X, 1 wash with milliQ-water. Colonies were counted under a stereo-microscope (NIKON-SMZ1500).

*1e/ 2i_OFF differentiation.*

Cells were adapted from FCS+LIF to 2i+LIF for 3-4 passages, harvested and seeded at ~5000-10000 cells/cm$^2$ in Poly-L-ornithine/Laminin coated cell culture treated surfaces, in 2i+LIF medium but ommiting PD0325901, CHIR99021 and LIF. The medium was changed every other day.

*1f/ Embryoid Bodies differentiation.*

Embryoid Bodies (EBs) imaged in Fig.S4A (top) were obtained with a hanging-drop protocol where a suspension of 0.1M cells/mL was distributed in 20 µl drops (2000 cells/drop) onto inverted cover plates of several dishes filled with PBS 1X to avoid evaporation. The day after, the aggregates formed in each drop were pooled and seeded on non-cell culture treated dishes and cultured in 10%FCS-DMEM without LIF for 8 days. EB differentiation for outgrowths (Fig.S4B, bottom) and RNA-seq analysis was performed by seeding 0.04M cells/$cm^2$. After 3 days, the aggregates were harvested (2min trypsinization) and transferred to non-cell culture treated dishes for 4 additional days in 10%FCS-DMEM medium without LIF. After 4 days, EBs were distributed at low density onto 0.1% gelatinized and cell culture treated dishes. They were kept in culture for 2 and 4 extra-days.

*1g/ Primitive endoderm differentiation.*

ES cells cultured in FCS+LIF were harvested and seeded onto 0.1% gelatinized single wells of µ-slide 4 well$^{Ph+}$ ibiTreat (Ibidi GmbH Ref#80446) at 37000 cells/$cm^2$ and differentiated as previously described (Anderson KGV et al., 2017). They were first cultured for 24H in endoderm base medium (EBM): RPMI 1640 Medium, GlutaMAX™ Supplement (Gibco, Cat# 61870036) supplemented with 2% B-27 minus insulin (Gibco, Cat# 15285074) and 100 µM 2-mercaptoethanol (Gibco, Cat# 31350-010). Subsequently, Activin A (20 ng/ml; R&D, Cat# 338-AC-010), CHIR99021 (3 µM) and LIF (10 ng/ml) were added (inductive PrE medium). The medium was changed every other day.

*1h/ Commitment assays.*

Cells cultured in 2i+LIF were plated at clonal density (600 cells per well of 6-well plates) and subject to 2i_OFF differentiation, as described above. After each day, the medium was changed into 2i+LIF for 7 days, after which cells were fixed and stained for alkaline phosphatase activity as described.

*1i/ Preparation of mitotic cells.*

To obtain mitotic ES cells (>95% purity as assessed by DAPI staining and microscopy), we used a nocodazole shake-off approach, as described before (Festuccia et al., 2018).

**2/ Flow cytometry activated cell sorting.**

Nanog-GFP cells (TNG; Chambers et al., 2017) were sorted using a Moflo Astrios with the "highest % of purity" parameter selected. After sorting, GFP-negative and GFP-positive sorted populations were reprocessed, with the same parameters used for sorting, to check the purity of each fraction (>95%).

**3/ Imaging.**

*3a/ Bright field microscopy.*

Cell culture pictures were taken on a Nikon Eclipse Ti-S inverted microscope equipped with: CFI S Plan Fluor ELWD ×20 objective; 89 North PhotoFluor LM-75; Hamamatsu ORCA-Flash 4.0LT camera; NIS Elements 4.3 software.

*3b/ Immunofluorescence of NANOG/OCT4 in E14Tg2a cells.*

Cells were trypsinized, counted and fixed for 10 min in 4% formaldehyde (Sigma Cat#F8775) at room temperature and quenched immediately after with Glycine 125 mM for 5 min at room temperature. After one wash in PBS 1X, they were cytospun at 0.5M cells/200 µl/spot (4 min – 300 rpm – low speed) on Superfrost+ slides. After washing the cells twice in PBS 1X, they were permeabilized with cold PBS 1X/0.5 % v/v Triton X-100 for 5 min, washed twice with cold PBS 1X, and blocked in PBS1X/1% Donkey Serum (DS) (Sigma, Cat# D9663) for 30 min on ice. Cells were incubated overnight at 4°C with primary antibodies (diluted in PBS1X/1% DS) within a humid chamber. After three washes in cold PBS1X, cells were incubated 1H at room temperature in the dark with secondary antibodies (diluted in PBS1X/1%DS), washed three times in PBS1X and nuclei counterstained with Vectashield antifade mounting medium with DAPI (Vectorlabs, Cat#H-1200). Imaging was performed with an inverted Nikon Eclipse X microscope equipped with: X20/0.45 (WD 8.2-6.9) objective; LUMENCOR excitation diodes; Hamamatsu ORCA-Flash 4.0LT camera; NIS Elements 4.3 software. Cell Profiler (Carpenter AE et al., 2006) was used for quantifications and ggplot2 (Wickham et al. 2016) for plotting in R.

*3c/ Immunofluorescence of NANOG in E14Tg2a cells and ΔK9 cells.*

WT and ΔK9 cells were trypsinized, counted and resuspended at 1M/ml in FCS free medium (DMEM-Glutamax/100 mM 2-mercaptoethanol/NEAA 1X) into sterile 1.5 mL Eppendorf tubes. Cells were then individually incubated either with 10 µM Rhodamine Red dye (InVitrogen, Cat#CMTPX C34552) or 1µM Deep Red dye (InVitrogen , Cat#C34565) for 30 min at 37°C. The labeled cells were then collected by centrifugation, washed with PBS1X, resuspended in DMEM/10%FCS+LIF medium and mixed at a 1:1 ratio for $WT^{Rhod}$ and $\Delta K9^{DeepRed}$ cells (usually ~0.4M each). The opposite labeling ($WT^{DeepRed}$ & $\Delta K9^{Rhod}$) was also performed with identical results. Mixed cells were seeded into a Poly-L-Ornithine/Laminin coated single well of a µ-slide 4 $well^{Ph+}$ ibiTreat (Ibidi GmbH Ref#80446) and incubated for ~6H at 37°C and 7% CO2. Cells were then fixed directly into the well by freshly prepared PFA 4% (Fisher Scientific, Cat# 16431755) for 10 min at room temperature in the dark, washed twice in PBS1X for 10 min and used immediately for immunostaining or stored at +4°C for short time. Cells were washed in PBS-Tw0.1% (PBS1X/0.1 % Tween20 (Sigma, Cat#P9416)) and permeabilised with PBS1X/0.1% Triton X-100 (Sigma, Cat#T8787) for 10 min at room temperature. After three washes with PBS-Tw0.1%, cells were blocked with PBS-Tw0.1%/10% Donkey Serum (Sigma, Cat#D9663) for 30 min on ice in the dark and incubated overnight with the primary antibody (diluted in PBS-Tw0.1%/10% DS). Following three washes with PBS-Tw0.1%, 1H incubation with secondary antibodies at room temperature in the dark and 2 washes with PBS1X, nuclei were counterstained with 4',6-diamidino-2-phenylindole (DAPI; Sigma, Cat# D9542), washed in PBS1X and use immediately for imaging or stored at +4°C for short time. Imaging was performed with an inverted Nikon Eclipse X microscope equipped with: X20/0.45 (WD 8.2-6.9) objective; LUMENCOR excitation diodes; Hamamatsu ORCA-Flash 4.0LT camera; NIS Elements 4.3 software. Quantifications were performed using Cell Profiler (Carpenter AE et al., 2006). For each experiment, E14Tg2a and ΔK9 cells quantifications were attributed using the FlowJo software. For each experiment, the fluorescence intensity of ΔK9 cells was normalised to the median of the corresponding E14Tg2a intensities imaged on the same spot. The data was ploted using the ggplot2 package (Wickham et al. 2016) in R.

*3d/ Immunofluorescence of NANOG and primitive endoderm markers.*

Differentiated cells were processed into their respective well: one wash with PBS1X, fixation with freshly prepared PFA 4% for 10 min at room temperature, three washes of PBS1X, and used immediately for immunostaining or stored at +4°C for short time. Immunostaining was performed as in 3c/ for NANOG in combination with primitive endoderm markers. Imaging was performed with an inverted Nikon Eclipse X microscope equipped with a x20 objective; LUMENCOR excitation diodes; Hamamatsu ORCA-Flash 4.0LT camera; NIS Elements 4.3 software. For each well, ~30 pictures were taken by randomly scanning the well. Quantifications were performed using Cell Profiler (Carpenter

AE et al., 2006) and ploted using the ggplot2 package (Wickham et al. 2016) in R. Representative images were generated with ImageJ software using identical settings for E14Tg2a and ΔK9 cells.

*3e/ Antibodies.*

Antibodies used for all immunostaining experiments are listed in Table S2.

**4/ Chromatin Immunoprecipitation.**

*4a/ Chromatin preparation.*

After trypsinisation, $10^7$ ES cells were crosslinked for 10 min in 3 ml DMEM/10% FCS/1% formaldehyde (Sigma Cat#F8775). Crosslinking was stopped with 125 mM glycine for 5 min at room temperature then 5min in ice. Cells were pelleted and washed with ice-cold PBS1X. Cells were resuspended in 1 ml of ice-cold swelling buffer (25 mM Hepes pH 7.95, 10 mM KCl, 10 mM EDTA) freshly supplemented with 1× protease inhibitor cocktail (PIC-Roche, Cat# 04 693 116 001) and 0.5% IGEPAL (Sigma, Cat#I8896). After 20 min on ice, the suspension was passed 50 times in a dounce homogenizer. Cells were then centrifuged and resuspended in 1 ml of ice-cold D3 (0.1% SDS, 15 mM Tris pH 7.6, 1 mM EDTA) buffer, freshly supplemented with 1× PIC. Samples were sonicated using a Covaris M220 - Setpoint @6°C and 10 cycles with the following parameters for each cycle : 60 sec duration, peak power of 67W, duty factor of 15% and cycles/burst of 500. A delay of 45 sec is added at the end of each cycle. Result of the average power is 10W. After centrifugation (15 min, 14000 rpm, 4 °C), the supernatant was stored at −80 °C until use. 20 µl were used to quantify the chromatin concentration and check DNA size (typically 200-600 bp).

*4b/ Immunoprecipitation.*

15 to 20 µg of chromatin were used for each ChIP after pre-clearing it for 1.5 hours rotating on-wheel at 4 °C in 1 ml of TSE150 (0.1% SDS, 1% Triton X-100, 2 mM EDTA, 20 mM Tris-HCl pH8, 150 mM NaCl) buffer containing 50 µl of protein G Sepharose beads (Active Motif, Cat#37499) 50% slurry, previously blocked with BSA (0.5 mg/ml ; Roche, Cat# 10711454001) and yeast tRNA (1 µg/ml ; Roche Cat# 10109495001). Immunoprecipitations were performed overnight rotating on-wheel at 4 °C in 500 µl of TSE150. 20 µl were set apart for input DNA extraction and precipitation. 50 µl of blocked protein G beads 50% slurry was added for 2 h rotating on-wheel at 4 °C. Beads were pelleted and washed for

5 min rotating on-wheel at room temperature with 1 ml of buffer in the following order: 2 × TSE150, 1 × TSE500 (as TSE150 but 500 mM NaCl), 1× washing buffer (10 mM Tris-HCl pH8, 0.25M LiCl, 0.5% IGEPAL, 0.5% Na-deoxycholate, 1 mM EDTA), and 2 × TE (10 mM Tris-HCl pH8, 1 mM EDTA). Elution was performed in 100 µl of elution buffer (1% SDS, 10 mM EDTA, 50 mM Tris-HCl pH 8) for 15 min at 65 °C after vigorous vortexing. Eluates were collected after centrifugation and beads rinsed in 150 µl of TE-1%SDS. After centrifugation, the supernatant was pooled with the corresponding first eluate. For both immunoprecipitated and input chromatin, the crosslinking was reversed overnight at 65 °C, followed by proteinase K treatment, phenol/chloroform extraction and ethanol precipitation.

*4c/ qPCR analysis.*

Input and IP samples were analysed by real-time quantitative PCR performed in duplicates in 384-well plates with a LightCycler 480 (Roche) using 4.6 µl of LightCycler 480 SYBR Green I Master (Roche, 04707516001), 5 µl of sample and 0.2 µl of each primer at 20 µM in a final reaction volume of 10 µl. Standard and melting curves were generated to verify the amplification efficiency (>85%) and the production of single DNA species. PCR primer sequences are listed in Table S2. The 2dCt method was used. All values were corrected to the input and ploted in R using ggplot2 (Wickham et al., 2016).

*4d/ Antibodies.*

Antibodies used for all ChIP experiments are listed in Table S2.

**5/ Gene expression analysis**

*5a/ RNA preparation, RT-qPCR and sequencing.*

E14Tg2a and ΔK9 cells were differentiated in parallel as embryoid bodies and cells recovered at d0, d4, d6, d8. RNA extraction and DNase treatment of 3 independent assays were made with NucleoSpin RNA Mini kit (Macherey Nagel, Cat# 740955.50) according to the manufacturer's protocol. Reverse Transcription was performed with 1 µg of total RNAs with random hexamers following manufacturer's protocol (Roche 04379012001); qPCR was performed as described above (*Tbp* was used as a reporter; see Table S2 for primer sequences). Stranded, poly-A selected RNA-seq libraries were prepared and sequenced (paired-end 150bp reads; around 50 millions each) by Novogene Co Ltd.

*5b/ Alignments and quantification.*

Stranded paired end RNA-seq reads were aligned to the mm10 genome using STAR (Dobin et al., 2013) and quantified by RSEM (Li and Dewey, 2011), with additional options " --calc-pme --calc-ci --estimate-rspd --forward-prob 0.0 --paired-end". Transcripts per million (TPM) were computed after omitting outliers identified by Principal Component Analysis (30 or more standard deviations away from the mean for PC1 component).

*5c/ Principal component analysis.*

PCA was run in R (prcomp function; with option center=TRUE) using log2 transformed TPM after selecting genes with at least 1 TPM in at least one sample mean (n=3). The top 1000 PC1 loadings (55% of variance) were used for gene ontology analyses. Data visualisation was made in base R.

*5d/ Differentially expressed genes.*

For all differential expression tests RSEM estimated read counts per sample were rounded for use with DESeq2 (Love et al., 2014), which was run without independent filtering. For differentially expressed genes in undifferentiated ES cells, we selected those with an FDR < 0.05 in ΔK9.1 versus E14Tg2a and ΔK9.2 versus E14Tg2a and same fold-change direction with abs(log2FC)>0.3. For embryoid body differentiation, we considered genes with absolute log2(FC) > 1 and FDR < 0.05 at any day of the differentiation versus undifferentiated cells for either E14Tg2a or ΔK9 cells. The corresponding heatmap was made in R with ComplexHeatmaps package (Gu et al., 2016).

*5e/ Clustering of differentially expressed genes and developmental annotation.*

K-means clustering was computed with R using the function kmeans with options k=6, nstart=50, iter.max=50. Only differentially expressed genes as identified during embryoid body differentiation were used (z scored mean TPM). The number of clusters was chosen as the minimal value identifying at least one cluster with maximal expression at each day of differentiation, including d0. Correlations to developmental gene expression were made by directly ploting the log2(FC) reported in a previous study using SCNMT-seq around gastrulation on mouse embryos (Argelaguet et al., 2019). The data was downloaded from ftp://ftp.ebi.ac.uk/pub/databases/scnmt_gastrulation. The heatmap was made in R with ComplexHeatmaps package (Gu et al., 2016) and the boxplot with ggplot2 (Wickham, 2016).

*5f/ Enrichment analyses.*

Gene Ontology analyses were performed using Enrichr (https://maayanlab.cloud/Enrichr/) and the top association was selected. To determine enrichments of each group of differentially expressed genes in proximity to TF binding sites for Nanog and Oct4/Sox2 (Festuccia et al., 2019), we calculated Fisher tests right tail p-values for the association between differentially expressed genes of each cluster within xbp of a binding site to a background of all genes clustered within xbp of a binding site, for x in [1, 1e+9] bp. Data visualization was made in base R.

### 6/ References.


Anderson KGV, Hamilton WB, Roske FV, Azad A, Knudsen TE, Canham MA, Forrester LM, Brickman JM. Insulin fine-tunes self-renewal pathways governing naive pluripotency and extra-embryonic endoderm. Nat Cell Biol. 2017 Oct;19(10):1164-1177. doi: 10.1038/ncb3617. Epub 2017 Sep 25. PMID: 28945231 DOI: 10.1038/ncb3617

Argelaguet R, Clark SJ, Mohammed H, Stapel LC, Krueger C, Kapourani CA, Imaz-Rosshandler I, Lohoff T, Xiang Y, Hanna CW, Smallwood S, Ibarra-Soria X, Buettner F, Sanguinetti G, Xie W, Krueger F, Göttgens B, Rugg-Gunn PJ, Kelsey G, Dean W, Nichols J, Stegle O, Marioni JC, Reik W. Multi-omics profiling of mouse gastrulation at single-cell resolution. Nature. 2019 Dec;576(7787):487-491. doi: 10.1038/s41586-019-1825-8. Epub 2019 Dec 11. PMID: 31827285; PMCID: PMC6924995.

Carpenter AE, Jones TR, Lamprecht MR, Clarke C, Kang IH, Friman O, Guertin DA, Chang JH, Lindquist RA, Moffat J, Golland P, Sabatini DM. CellProfiler: image analysis software for identifying and quantifying cell phenotypes. Genome Biol. 2006;7(10):R100. doi: 10.1186/gb-2006-7-10-r100. Epub 2006 Oct 31. PMID: 17076895; PMCID: PMC1794559.

Chambers I, Silva J, Colby D, Nichols J, Nijmeijer B, Robertson M, Vrana J, Jones K, Grotewold L, Smith A. Nanog safeguards pluripotency and mediates germline development. Nature. 2007 Dec 20;450(7173):1230-4. doi: 10.1038/nature06403. PMID: 18097409.

Festuccia N, Owens N, Papadopoulou T, Gonzalez I, Tachtsidi A, Vandoermel-Pournin S, Gallego E, Gutierrez N, Dubois A, Cohen-Tannoudji M, Navarro P. Transcription factor activity and nucleosome organization in mitosis. Genome Res. 2019 Feb;29(2):250-260. doi: 10.1101/gr.243048.118. Epub 2019 Jan 17. PMID: 30655337; PMCID: PMC6360816.

Gu Z, Eils R, Schlesner M. Complex heatmaps reveal patterns and correlations in multidimensional genomic data. Bioinformatics. 2016 Sep 15;32(18):2847-9. doi: 10.1093/bioinformatics/btw313. Epub 2016 May 20. PMID: 27207943.

Heurtier V, Owens N, Gonzalez I, Mueller F, Proux C, Mornico D, Clerc P, Dubois A & Navarro P.The molecular logic of Nanog-induced self-renewal in mouse embryonic stem cells
Nature Communications volume 10, Article number: 1109 (2019) doi: 10.1038/s41467-019-09041-z. PMID: 30846691

Kunath T, Arnaud D, Uy GD et al. Imprinted X-inactivation in extra-embryonic endoderm cell lines from mouse blastocysts. Development 2005;132:1649–1661.

Tanaka S, Kunath T, Hadjantonakis AK et al. Promotion of trophoblast stem cell proliferation by FGF4. Science 1998;282:2072– 2075.

Wickham, H. (2016). ggplot2: Elegant Graphics for Data Analysis. Springer-Verlag New York.
